# Heavy metal sensitivities of gene deletion strains for *ITT1* and *RPS1A* connect their activities to the expression of *URE2*, a key gene involved in metal detoxification in yeast

**DOI:** 10.1101/331009

**Authors:** Houman Moteshareie, Maryam Hajikarimlou, Alex Mulet Indrayanti, Daniel Burnside, Ana Paula Dias, Clara Lettl, Duale Ahmed, Katayoun Omidi, Tom Kazmirchuk, Nathalie Puchacz, Narges Zare, Sarah Takallou, Thet Naing, Raúl Bonne Hernández, William G. Willmore, Mohan Babu, Bruce McKay, Bahram Samanfar, Martin Holcik, Ashkan Golshani

## Abstract

Heavy metal and metalloid contaminations are among the most concerning types of pollutant in the environment. Consequently, it is important to investigate the molecular mechanisms of cellular responses and detoxification pathways for these compounds in living organisms. To date, a number of genes have been linked to the detoxification process. The expression of these genes can be controlled at both transcriptional and translational levels. In baker’s yeast, *Saccharomyces cerevisiae*, resistance to a wide range of toxic metals is regulated by glutathione S-transferases. Yeast *URE2* encodes for a protein that has glutathione peroxidase activity and is homologous to mammalian glutathione S-transferases. The *URE2* expression is critical to cell survival under heavy metal stress. Here, we report on the finding of two genes, *ITT1*, an inhibitor of translation termination, and *RPS1A*, a small ribosomal protein, that when deleted yeast cells exhibit similar metal sensitivity phenotypes to gene deletion strain for *URE2*. Neither of these genes were previously linked to metal toxicity. Our gene expression analysis illustrates that these two genes affect *URE2* mRNA expression at the level of translation.

**Summary statement:** We identified two yeast genes, *ITT1* and *RPS1A*, that when deleted, results in yeast cells sensitivity to heavy metals and metalloids. Further investigation indicated that they influence the expression of *URE2* gene, a key player in metal detoxification, by upregulating its translation. Our findings suggest that *ITT1* and *RPS1A* play an indirect role in responding to toxic metal stress.

## Introduction

Heavy metals and metalloids comprise a group of elements that are loosely defined as relatively high-density transition metals and metalloids [1], [2]. Different metals are found in varied concentrations across the environment. Some of these heavy elements, such as iron (Fe), cobalt (Co) and zinc Zn, are essential nutrients, while others are relatively harmless at low concentrations such as rubidium (Ru), silver (Ag) and indium (In). At higher concentrations, all metals and metalloids derived from natural environment [3] or anthropogenic sources such as phosphate fertilizers, disinfectants, fungicides, sewage sludge, industrial waste, bad watering practices in agricultural lands, and dust from smelters [4], [5] are toxic to living cells [6], [7], [8]. Among these, arsenic (As) is one of the most toxic despite being the twentieth most abundant element on our planet. Its inorganic oxyanion forms including arsenite (As(III)) and arsenate (As(II)) are highly lethal to living organisms [9].

Over the course of evolution, many organisms have found ways, for example, by evolving molecular pathways to survive increased concentrations of metallic toxins in their environment e.g. [10], [11], [12], [13]. Microbes with extreme adaptation to heavy metals use detoxification pathways to reduce toxic metals to a lower redox state, which lessens their mobility and toxicity [14]. The baker’s yeast, *Saccharomyces cerevisiae* possesses an effective mechanism to negate heavy metal and metalloids toxicity, allowing it to survive a broad range of toxic stress scenarios [15], [16]. This makes yeast an ideal model organism to study molecular mechanisms of the stress response that drive detoxification processes.

The glutathione S-transferases (GSTs) are key enzymes that mediate the resistance of *S. cerevisiae* to a wide range of heavy metals and metalloids. Yeast *Ureidosuccinate Transport 2* (*URE2*) gene product is structurally homologous to mammalian GST and is a major player in the detoxification of *S. cerevisiae* against toxic metals through its glutathione peroxidase activity [17], [18]. For detoxification purposes, GST proteins catalyze the conjugation of the reduced form of glutathione (GSH) to xenobiotic substrates [19]. The deletion strain for *URE2* is hypersensitive to a wide range of heavy metals and metalloids including As, Cd and nickel (Ni) [19], [20]. In this report, we show that the deletion of either *ITT1* (inhibitor of translation termination 1) or *RPS1A* (small ribosomal subunit protein 10), makes the cells more sensitive to As(III), cadmium (Cd) and Ni suggesting a functional connection of these two genes with heavy metal toxicity. Itt1p is known to modulate the efficiency of translation termination through physical interactions with two eukaryotic release factors eRF1 (Sup45p) and eRF3 (Sup35p) [21]. Rps1Ap is a constituent of the small ribosomal subunit; little information is known about its function in *S. cerevisiae* [22]. Neither of these genes had previously been linked to heavy metal toxicity. Overall, we provide evidence that the connection for *ITT1* and *RPS1A* with heavy metal toxicity is through their influence on the translation of *URE2* gene.

## Materials and methods

### Strains and plasmids used in this study

Yeast, *S. cerevisiae*, mating type (a) MATa strain Y4741 (*MATa orfΔ::KanMAX4 his3Δ1 leu2Δ0 met15Δ0*) and mating type (α) MAT*α* strain, BY7092 (*MATα Can1Δ::STE2pr-HIS3 Lyp11Δ leu21Δ0 his31Δ0 met151Δ0*) [23] were utilized for this study. The yeast MATa knockout (YKO) collection [23] and PCR-based transformed cells were used as a source of gene deletion mutants; the open reading frame (ORF) collection [24] was used for over expression plasmid vectors. Yeast GFP Clones [25] were modified for Western blot analysis. pAG25 plasmid containing the Nourseothricin Sulfate (clonNAT) resistance gene was used as a DNA template in PCR to generate gene knockouts. *Escherichia coli* strain *DH5α* was used to replicate different plasmids [26]. Two plasmids were used carrying the *β-galactosidase* open reading frame for quantification of *URE2*-IRES and cap-dependent translation activities. p281-4-*URE2* contained a *URE2*-IRES region which was fused with *β-galactosidase* gene, and p281 contained only the *β- galactosidase* gene as a control for cap-dependent translation [27]. All plasmids carried an ampicillin resistance gene which was used as selectable marker in *E. coli* and the *URAcil requiring 3* (*URA3*) marker gene which was used for selection in yeast.

### Media and MiniPrep

YP (1% Yeast extract, 2% Peptone) or SC (Synthetic Complete) with selective amino acids (0.67% Yeast nitrogen base w/o amino acids, 0.2% Dropout mix,) either with 2% dextrose or 2% galactose as a source of carbohydrates was used as culture medium for yeast and LB (Lysogeny Broth) was used for *E. coli* cultures. 2% agar was used for all solid media. Yeast cells were grown at 30°C unless otherwise indicated in the SGA procedure. *E. coli* cells were grown at 37°C. Yeast plasmid extraction was performed by using quick plasmid kit E.Z.N.A. Yeast plasmid mini kit (Omega Bio-tek®) and *E. coli* plasmid extraction was carried out by using GeneJET plasmid miniprep kit (Thermofisher®) according to the manufacturer’s instructions.

### Gene knockout and DNA transformation

Mutant strains were either selected from the library of gene deletions [23] in MATa haploid form or a PCR-based gene knockout approach was used to achieve gene deletion. Targeted gene knockout strains were generated by PCR-based gene deletion strategy utilizing the clonNAT selection gene [28], [29]. Plasmid and gene transformation were performed by using a chemical-transformation strategy (LiOAc method) and confirmed via colony PCR [30], [31].

### Chemical sensitivity

Colony count assay (spot test) was performed to estimate the number of viable cells, based on their ability to give rise to colonies. Strains were grown in liquid YPD or SC without uracil to saturation phase and serially diluted in dH_2_O to 10^−4^ and aliquots were streaked on solid media. The cells were cultured on YP, and YP supplemented with Na_3_AsO_3_ (As(III) (1 mM)), CdCl_2_ (Cd (0.1 mM)), NiCl_2_ (Ni (8 mM)), as well as 6% ethanol and 6% ethanol + Na3AsO_3_ (As(III) (0.3 mM)) for two days at 30°C. YP + 2% glucose (YPD) was used for the experiments without the involvement of any plasmid and YP + 2% galactose (YPG) was used for experiments harboring plasmids (overexpression) with a galactose-inducible promoter (GAL1/10), in order to activate the desired gene. The number of colonies under drug conditions were compared to the number of colonies formed under non-drug conditions for consistency and subsequently normalized to the number of colonies formed by wild type (WT) strain under the same condition. Each experiment was repeated at least three times.

### Quantitative β-galactosidase assay

*ortho*-Nitrophenyl-β-galactoside (ONPG)-based *β-galactosidase* analysis was used to quantify involvement of the identified genes in the IRES-mediated translation of *URE2* via quantification of β-galactosidase activity produced by a plasmid containing *URE2*-IRES fused to a *β-galactosidase* reporter (p281-4-*URE2*) [27]. This plasmid contains a DNA sequence that forms four strong hairpin loops prior to *URE2*-IRES region at its mRNA level. These four hairpin loops inhibit cap-dependent translation of *URE2* mRNA. A background plasmid (p281) carrying only a *β-galactosidase* reporter was used as a control for cap-dependent translation [27]. Both plasmids contained a GAL1/10 promoter and YPG was used as a medium to activate the desired gene [32]. Each experiment was repeated at least three times.

### Reverse Transcriptase quantitative PCR (RT-qPCR)

This methodology was used to assess the content of target mRNAs. Total mRNA was reverse-transcribed into complementary DNA (cDNA), using iScript Select cDNA Synthesis Kit (Bio-Rad®) according to the manufacturer’s instructions. cDNA was then used as a template for quantitative PCR (qPCR). Total RNA extractions were performed with Qiagen RNeasy Mini Kit (Qiagen®). qPCR was carried out using Bio-Rad iQ SYBR Green Supermix and the CFX connect real time system (Bio-Rad®), according to the manufacturer’s instructions. In this experiment, *PGK1* was used as a constitutive housekeeping gene and related to WT [31], [32]. Each qPCR experiment was repeated at least three times using separate cDNA samples.

### Immunoblotting

Western blotting was used to quantify relative protein levels. Total protein was isolated using detergent-free methods as described in [33]. Samples were grown overnight in liquid YPD, pelleted and washed with PBS buffer. Samples for As(III) treatment, were grown overnight in liquid YPD and then treated with Na_3_AsO_3_ (As(III) (0.5 mM)) for 2 hours. Bicinchoninic acid assay (BCA) (Thermofisher®) was used to quantify total protein concentration according to manufacturer’s instructions. Equal amounts of protein were loaded onto a 10% SDS-PAGE gel, run on Mini-PROTEAN Tetra cell electrophoresis apparatus system (Bio-Rad®) [34]. Proteins were transferred to a nitrocellulose 0.45 μm paper (Bio-Rad®) via a Trans-Blot Semi-Dry Transfer (Bio-Rad®). Mouse monoclonal anti-GFP antibody (Santa Cruz®) was used to detect protein level of Ure2p in Ure2-GFP protein fusion strains. Pgk1p was also used as a constitutive housekeeping protein for quantification purposes. Mouse anti-Pgk1 (Abcam®) was used to detect Pgk1p levels [31]. Immunoblots were visualized with chemiluminescent substrates (Bio-Rad®) on a Vilber Lourmat gel doc Fusion FX5-XT (Vilber®). Densitometry analysis was carried out using the FUSION FX software (Vilber®), and each density was normalized to the density formed by Pgk1p control. Each experiment was repeated at least three times using three separate total protein isolates.

### Polyribosome fractionation

The total RNA extraction from yeast cells was adapted from [35]. Overnight cell culture was used to inoculate YPD liquid medium. Prior to extraction, cycloheximide (100 μg/ml was added to the samples for 15 minutes. Yeast culture was harvested at mid-log phase (OD_600_ 0.6– 0.8) and immediately chilled on dry ice prior to lysing the cells. Yeast cells were lysed by mechanical disruption using 425 – 600 μm acid-washed glass beads (Sigma®). Fifty μg of the total RNA was then loaded on a 10–50% sucrose gradient (20 mM Tris pH 8, 140 mM KCl, 5 mM MgCl_2_, 0.5 mM DTT, 100 μg/ml cycloheximide and sucrose to desired concentration). An Automated Gradient Maker (Biocomp gradient maker) was used to produce the sucrose gradients. Centrifugation was performed at 40,000 rpm for 2 hours at 4°C (Beckman Optima LE-80K Ultracentrifuge) to separate the particles according to relative density.

Samples were analysed via a Biocomp Gradient Station immediately after centrifugation. The instrument recorded A_254_ using a flow cell coupled with a spectrophotometer (Bio-Rad Econo UV monitor). In this procedure, untranslated mRNAs (top fractions) are separated from polysome-associated mRNAs (bottom fractions) as described in [36]. Fractions were collected (~650 μl) using Bio-Rad Collection Station (Bio-Rad®) and adjusted to 1% SDS for fluctuation analysis of *URE2* mRNA level via RT-qPCR [37]. Each polysome profiling experiment was repeated at least three times.

Luciferase RNA (0.1 μg/ml) (Promega®) was then added to each fraction as a control. RNA was precipitated overnight and purified as described in [37], by using Glycoblue and acidic Phenol/chloroform (pH 4). Purified RNA samples were subjected to quantification for *URE2* mRNA by performing RT-qPCR as described in the above section. The normalized values for each fraction was determined by using the Cq values for *URE2* and the Cq values for the gene of interest using the formula [2λ (Cq_luciferase_ – Cq_target gene_)] [38]. The relative amount of *URE2* mRNA was calculated by dividing the amount in each fraction by the total signal in all fractions [38], [39]. RT-qPCR analysis for polysome profiling was repeated at least three times using fractions from three separate polysome profiles.

### Genetic interaction (GI) and conditional GI analysis

Synthetic genetic array (SGA) analysis was performed in large-scale through the creation of double-mutations and subsequent analysis of colony size (fitness) as previously described in [31], [40]. In summary, both gene candidates *ITT1* and *RPS1A* were knocked out in MATα (BY7092) and crossed with two arrays of haploid MATa knock-out strains [31]. The first array contained 384 deletion strains for genes that are directly or indirectly involved in the process of translation. The second contained 384 random genes that were selected from the YKO collection [23], [31], which was used as a control. Selectable markers designated in each background mating type, allowed for multiple selection steps. Meiotic progeny harboring both mutations were selected. The created arrays could then be used to score double mutants for their altered fitness under certain conditions [23].

Colony size of both single mutant arrays (reference and control) and double mutant arrays were measured for their colony fitness [41], [42]. After three repeats, the interactions with 20% alteration or more in at least two repeats were considered positive hits. Conditional SGA analysis was carried on under sub-inhibitory concentrations of chemicals (0.7 mM for As(III) and 60 ng/ml for cycloheximide). PSA analysis was performed under a high sub-inhibitory targeted condition as previously described in [31], [43] (As(III) (1.2 mM) and cycloheximide (100 ng/ml)). Each experiment was repeated three times and the interactions with 20% alteration or more in at least two screens were scored as positive.

## Results and discussion

### Deletion of *ITT1* or *RPS1A* increases yeast sensitivity to heavy metals

Understanding the biology of the stress that heavy metals exert on a cell, as well as the cellular responses and mechanisms that a cell uses for detoxification of these toxins, has been the subject of numerous investigations over the past decades e.g. [2], [14], [15], [16]. Although much has been learned, additional studies are needed to uncover details of such responses as well as additional genes that may participate in this process. To this end, while screening for yeast gene deletion mutants against heavy metals, we identified two deletion mutant strains for *ITT1* and *RPS1A* that showed increased sensitivity to three heavy metals (As, Ni and Cd). In our spot test sensitivity analysis, when As(III) (1 mM), Cd (0.1 mM) and Ni (8 mM) were added to the solid media the number of normalized yeast colony counts were significantly reduced for *Δitt1* and *Δrps1a* strains (Fig 1a), highlighting the sensitivity of the mutant strains to these metals and metalloids. Rescue experiments revealed a complete recovery to heavy metal toxicity upon the reintroduction of the deleted genes into their corresponding mutants (Fig1b). This reversion of sensitivity indicates that the sensitivity phenotypes are in fact a consequence of the intended gene deletions and are not due to a possible secondary mutation within the genome. The sensitivity of deletion mutants to As, Cd and Ni suggests a potential association for the target genes to heavy metal toxicity, a unique observation that has not been previously reported.

**Figure 1:**
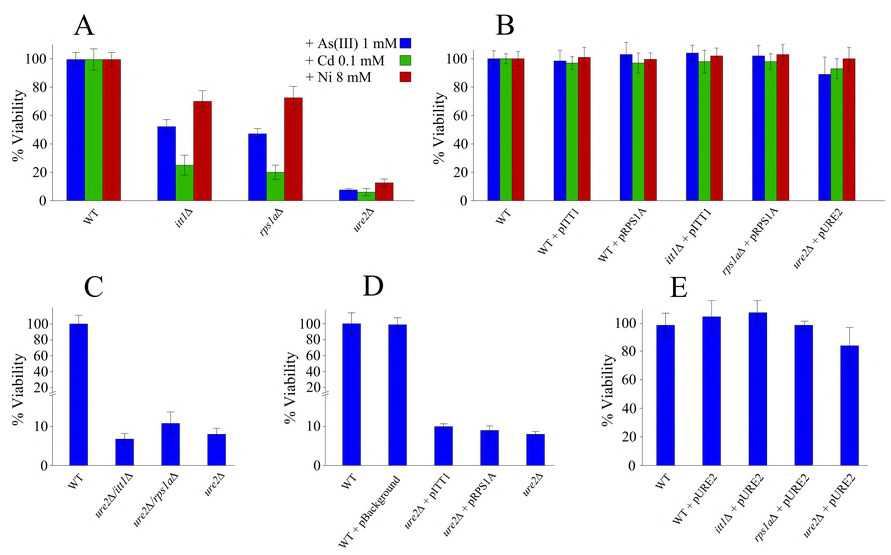
Normalized CFU counts for different yeast strains after exposure to As(III) (1 mM). CFU counts after exposure to the experimental condition are normalized to control condition counts and expressed as a percentage of the average CFU counts. (A) Sensitivity of *itt1Δ, rps1aΔ* and *ure2Δ* compare to WT phenotype. (B) Rescued sensitivity of all deletion strains by reintroduction of their overexpression plasmids. (C) Sensitivity analysis for double gene deletions for *ITT1* or *RPS1A* in absence of *URE2* compared to single gene deletion of *URE2.* (D) Sensitivity analysis for overexpression of *ITT1* or *RPS1A* in absence of *URE2*. (E) sensitivity of *itt1Δ, rps1aΔ* after introduction of pURE2 (carries *URE2* genes). Each experiment was repeated at least three times. Error bars are calculated as standard deviations. Colour code is the same as in (A) for all panels.

Since the function of both *ITT1* and *RPS1A* can be linked to the process of protein biosynthesis [21], [22], it is conceivable that these genes may indirectly influence heavy metal sensitivity by regulating the activity of another gene. However, given that the Ure2p is reported to be a key enzyme involved in heavy metal detoxification in yeast [19], [20], we made double gene deletion mutants for *ITT1* and *RPS1A* with *URE2* and exposed them to As(III) (1 mM) for further analysis (Fig 1c). The sensitivity analysis of the double gene deletion mutants indicated no increased sensitivity to As(III) in addition to that observed for the single gene deletion mutant for *URE2*. This specifies a dominant effect for *URE2* on heavy metal sensitivity over *ITT1* and *RPS1A* (Fig 1c). To further support the observed phenotype, we introduced plasmid vectors containing *ITT1* and *RPS1A* into the deletion strain of *URE2* (Fig 1d). The results demonstrated the same levels of sensitivity to As(III) with no compensation, deeming indirect roles of *ITT1* and *RPS1A* in rescuing the cells from As(III) toxicity when *URE2* is deleted. One way to explain this data is that *ITT1* and *RPS1A* may exert their effect on sensitivity via the same pathway as *URE2*. If *ITT1* and *RPS1A* influenced a second pathway, it might be expected that their deletion would have had an additional effect on sensitivity when combined with *URE2* deletion [44]. However, this was not observed. On the other hand, overexpression of *URE2* in *itt1Δ and rps1aΔ* strains reversed the As(III) sensitivity phenotype observed by the corresponding gene deletions, effectively deeming these gene deletions inconsequential for heavy metal sensitivity (Fig 1e). These observations are in accordance with the activity of *URE2* as a dominant player in heavy metal toxicity and that it functions downstream of *ITT1* and *RPS1A.*

Our findings suggest that the influence of *ITT1* and *RPS1A* on heavy metal sensitivity is linked to *URE2*. Although both *ITT1* and *RPS1A* have reported roles in protein biosynthesis [21], [22], the possible mechanism, regulation of transcription or translation of *URE2* remains to be investigated.

### *ITT1* and *RPS1A* do not affect the expression of *URE2* at the mRNA level

RT-qPCR was employed to detect possible changes to *URE2* mRNA levels in the absence of *ITT1* and *RPS1A*. Figure 2 illustrates our observation that in comparison to WT, the *URE2* mRNA appeared unchanged in the mutant strains, *Δitt1* and *Δrps1a*. We also investigated the content of *URE2* mRNA after exposure to As(III) (0.5 mM). As noted before, neither the deletion of *ITT1* or *RPS1A* appeared to influence the content of *URE2* mRNA, suggesting that the activity of *ITT1* and *RPS1A* is independent of *URE2* mRNA content.

**Figure 2:**
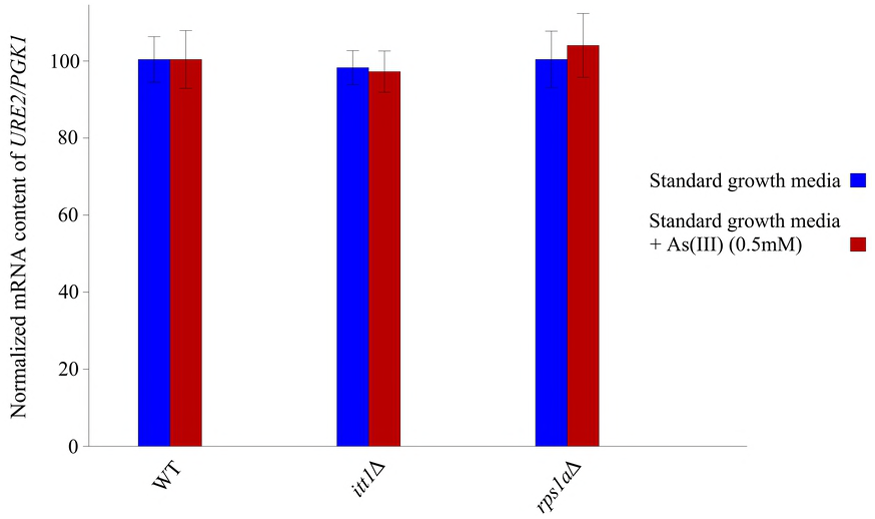
The relative *URE2* mRNA level quantified by normalizing the mRNA content of the mutant strains to those in the wild type. The house keeping gene *PGK1* was used as an internal control. Deletion of *ITT1* or *RPS1A* had no effect on the normalized *URE2* mRNA content. Each experiment was repeated at least three times. Error bars represent standard deviations.

### Ure2p content is reduced in the absence of *ITT1* or *RPS1A*

We investigated the level of Ure2p in the presence and absence of *ITT1* or *RPS1A* in the cells by Western blot analysis. This was accomplished using a strain that carried Ure2p fused to a GFP at the genomic level. Our analyses show a reduction in the endogenously-expressed Ure2-GFP fusion protein levels in the absence of *ITT1* or *RPS1A* (Fig 3). When either *ITT1* or *RPS1A* were deleted, Ure2p levels were reduced by approximately 40% and 60%, respectively, compared to the WT cells (Fig 3a). This suggests that *ITT1* and *RPS1A* play an imperative role in regulating the expression of Ure2p. In parallel, introduction of As(III) (0.5 mM) to the growth media reduced Ure2p approximately 70% and 80% for *Δitt1* and *Δrps1a,* respectively compared to WT strain (Fig 3b). Deletion of *ITT1* or *RPS1A* did not change the protein levels of Pgk1p, used as an internal control. These observations provide evidence by connecting *ITT1* and *RPS1A* activities to the level of Ure2p. The additional reduction in the Ure2p levels in the presence of As(III) implicates that *ITT1* and *RPS1A* may have a higher influence in regulation of Ure2p expression under a stress condition.

**Figure 3:**
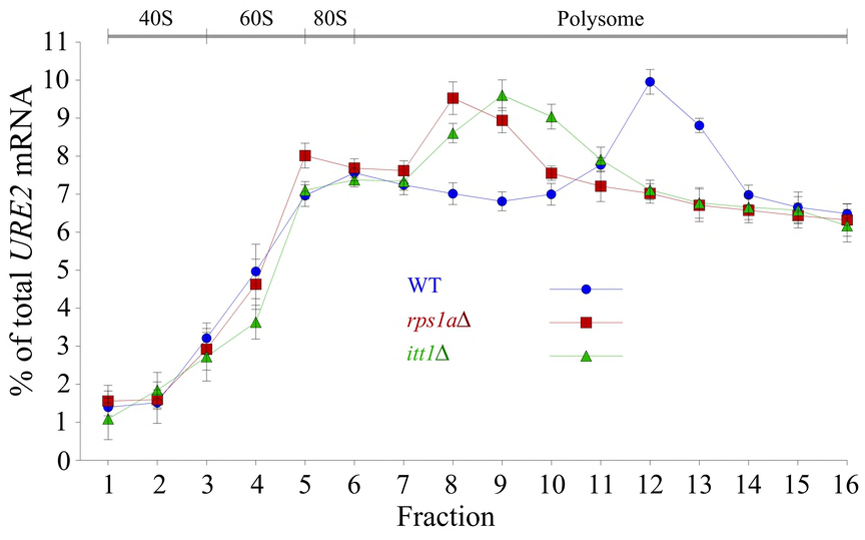
Western blot followed by densitometry analysis to measure Ure2-GFP levels in different yeast strains. Values are normalized to that for Pgk1p, used as an internal control and related to the values for WT strain. (A) Cells are grown under standard laboratory conditions. (B) Cells are challenged by As(III) (0.5 mM). Each experiment was repeated at least three times. Error bars represent standard deviations.

### Analysis of the URE2 translation

Since our data suggests a role for *ITT1* and *RPS1A* in modulating *URE2* expression at the protein level, polyribosome-bound mRNA analysis was performed. In this method, fractions of polysomes are isolated and analyzed for their content of a target mRNA. Those mRNAs that are translated more efficiently are generally found in association with multiple ribosomes and hence will be isolated in heavier polysome fractions. In contrast, those mRNAs, which are translated to a lesser degree, can be found in lower density fractions [34]. In this way, the distribution of mRNAs within polysome fractions can be used to estimate the translation efficiency of the target mRNA. Using this strategy, polysome profile analysis was performed for *URE2* mRNA, in the presence or absence of *ITT1* and *RPS1A*. Analysis of the polysome fractions for *URE2* mRNA content using RT-qPCR, normalized to a control (housekeeping) mRNA (*PGK1*), showed a shift in *URE2* mRNA accumulation towards lighter polysome fractions for mutant strains (Fig 4), suggesting that when *ITT1* or *RPS1A* are deleted, *URE2* mRNA is translated to a less efficiently. These data provide direct evidence that *ITT1* and *RPS1A* affect the translation of *URE2* mRNA.

**Figure 4:**
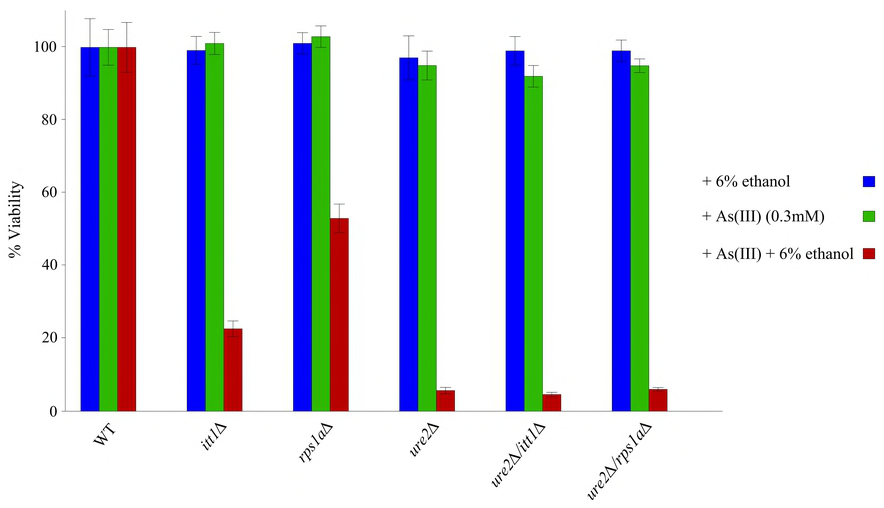
Polysome-bound mRNA analysis of *Δitt1, Δrps1a* and WT strains. The amount of *URE2* mRNA in each fraction was determined by RT-qPCR and the percentage of total *URE2* mRNA on the gradient is plotted for each fraction. The profiles of *PGK1* mRNA, used as an internal control, were similar for deletion and WT strains and were used to normalize other values. Each experiment was repeated at least three times. Error bars represent standard deviations.

### Ethanol increases As(III) sensitivity for *ITT1* and *RPS1A* gene deletion strains

In addition to cap-dependent translation, *URE2* mRNA has been shown to undergo a cap-independent translation, which represents an interesting mode of gene expression control [27]. Cap-dependent translation is mediated through the scanning of mRNA 5’ UTR to find a suitable start codon. In cap-independent translation, mRNA structures called Internal Ribosome Entry Site (IRES) mediate the interaction between ribosomes and the mRNA, independently of the 5’cap [45]. IRES-mediated translation is mainly used by RNA viruses but it can also be found in cellular mRNAs [46], [47]. The majority of translation in eukaryotes occurs through cap-dependent translation, whereas IRES-mediated translation is often associated with physiological conditions such as stress, where general translation is compromised [45], [48], [49].

Knowing that the exposure to heavy metal causes a stress condition for yeast cells e.g. [19], [50] we examined the possibility that *ITT1* and *RPS1A* may influence IRES-mediated *URE2* mRNA translation. As a quick measure to examine this possibility, ethanol sensitivity analysis was used as a positive stress control. Eukaryotic initiation factor 2A (eIF2A) acts as a regulator of IRES-mediated translation in *S. cerevisiae* cells [51]. Abundance of eIF2A is shown to specifically repress IRES-mediated translation of *URE2* as well as other yeast genes with IRES forming region [51]. It was previously reported that ethanol reduces the expression of eIF2A at protein levels and hence promotes translation via IRES elements [51].

As described, *URE2* plays a critical role in heavy metal detoxification [19], [49]. If *ITT1* and *RPS1A* affect IRES-mediated translation of *URE2* mRNA, one may expect the presence of ethanol to further enhance the sensitivity of *itt1Δ and rps1aΔ* to As(III) treatment. In this case, presence of ethanol and deletion of either *ITT1* or *RPS1A* could be considered to introduce a double effect on the same overall process.

As demonstrated in Figure 5, *ure2Δ, itt1Δ* and *rps1aΔ* showed hypersensitivity to a very low concentration of As(III) (0.3 mM) in the presence of 6% ethanol which suggests that ethanol increases the sensitivity of mutant strains to As(III). The mutant strains did not exhibit sensitivity to neither 0.3 mM of As(III) nor 6% ethanol separately. These data suggest a connection for the activities of *ITT1* and *RPS1A* to IRES-mediated translation. As well, a double gene deletion for *ITT1* or *RPS1A* with *URE2* did not result in additional sensitivity compared to that observed for *URE2* single gene deletion. This additionally reiterates the connection between the activities of *ITT1* and *RPS1A* via *URE2* expression.

**Figure 5:**
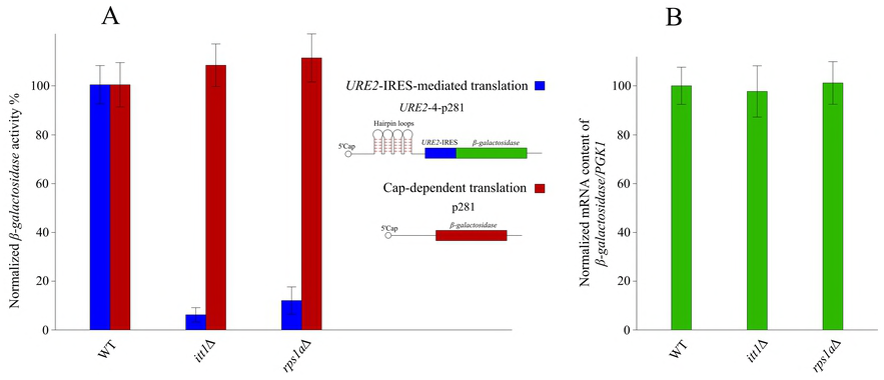
Average viability of single and double gene deletion strains for *ITT1* and *RPS1A* with *URE2.* Cells were exposed to As(III) (0.3 mM), 6% ethanol, a combination of both (6% ethanol + As(III) (0.3 mM)) or no treatment (control). CFU counts after exposure to the experimental conditions are normalized to CFU counts for WT strain and expressed as a percentage of the average CFU counts. Error bars represent standard deviation of at least three independent experiments.

### *ITT1* and *RPS1A* affect IRES-mediated translation of a *β-galactosidase* reporter gene

To further study the effect of *ITT1* and *RPS1A* on IRES-mediated translation of *URE2*, we used a *β-galactosidase* reporter whereby the *β-gal* mRNA is under the translational control of the *URE2*-IRES element. For this purpose, we utilized a previously published plasmid construct known as p281-4-*URE2* (Fig 6) [27], [51].

**Figure 6:**
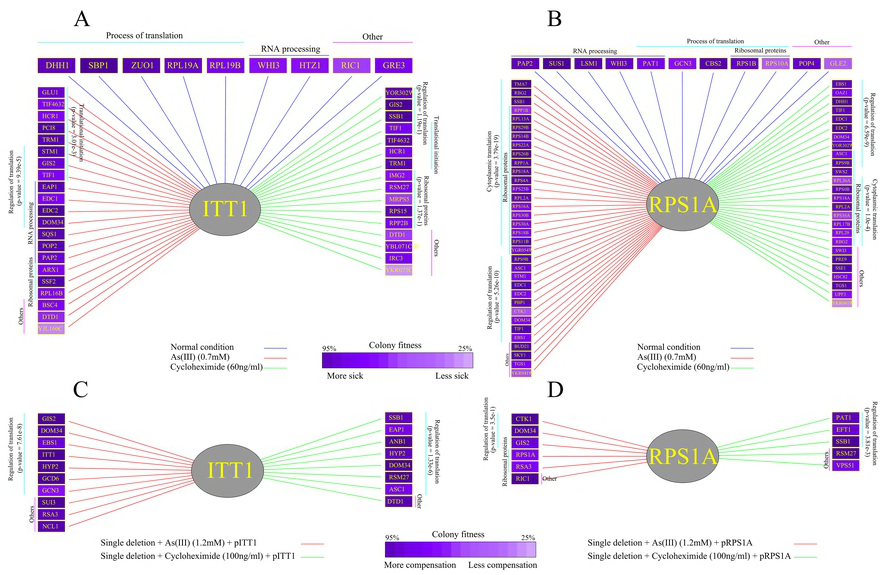
Effect of gene deletion on translation and transcription of *β-galactosidase* mRNA. (A) The relative β-galactosidase activity is determined by normalizing the activity of the mutant strains to that of the WT strain. Blue bars represent β-galactosidase activity under the translational control of *URE2*-IRES. Red bars represent β-galactosidase activity via cap-dependent translation. (B) The relative *β-galactosidase* mRNA level quantified by normalizing the mRNA content of the mutant strains to those in the wild type. The house keeping gene *PGK1* was used as an internal control. Deletion of *ITT1* or *RPS1A* had no effect on the normalized *β-galactosidase* mRNA content. Each experiment was repeated at least three times. Error bars represent standard deviations.

As shown in Figure 6, deletion of either *ITT1* or *RPS1A* resulted in reduced expression of *β-galactosidase* under the translational control of *URE2*-IRES. This data further supports the notion that *ITT1* and *RPS1A* regulate *URE2*-IRES-mediated translation. As a control to account for cap-dependent translation activity, p281 background construct [27], [51] carrying a *β-galactosidase* mRNA lacking four hairpin loops and *URE2*-IRES was utilized. We observed no significant difference in *β-galactosidase* activity for WT, *Δitt1* and *Δrps1a* strains indicating that *ITT1* and *RPS1A* do not influence cap-dependent translation (Fig 6). Since the reduced levels of β-galactosidase may be a consequence of reduced mRNA, the *β-galactosidase* mRNA was evaluated using RT-qPCR. As expected, we observed no significant change in its mRNA level in the absence of neither *ITT1* nor *RPS1A* (Fig 6b).

### Genetic interaction analysis further connects the activity of *ITT1* and *RPS1A* to regulation of translation in response to stress

To further examine the role of *ITT1* and *RPS1A* in the process of translation, we studied the genetic interactions (GIs) by screening *ITT1* and *RPS1A* against an array of 384 genes associated with protein biosynthesis and a second set of 384 random genes used for control purposes. Genes that are functionally related often partake in GIs (also known as epistatic interactions) [52], [53]. The most commonly studied form of GI is known as negative GI, where the reduced fitness or lethal phenotype of a double mutant strain for missing two genes is not observed in single mutant strains [31]. Genes that are associated with parallel and compensating pathways are thought to commonly form negative GIs [31], [40]. Under standard laboratory growth conditions, both *ITT1* and *RPS1A* exhibited negative GIs with a limited number of genes involved in the process of protein biosynthesis (Figs 7a and b). This is expected as both genes are thought to be involved in the process of translation. Examples of genes that formed GIs with *ITT1* and *RPS1A* are large ribosomal subunit protein 19A (*RPL19A*) and general control non-derepressible (*GCN3*), respectively. Rpl19p is a conserved large ribosomal subunit protein involved in ribosomal intersubunit bridging and its alteration is connected to the fidelity of translation [54], [55]. On the other hand, *GCN3* is the alpha subunit of translation initiation factor 2B (eIF2B), which is involved in guanine-nucleotide exchange for eIF2. In this manner, phosphorylation of eIF2B regulates the activity of Gcn3p [56], [57].

**Figure 7:**
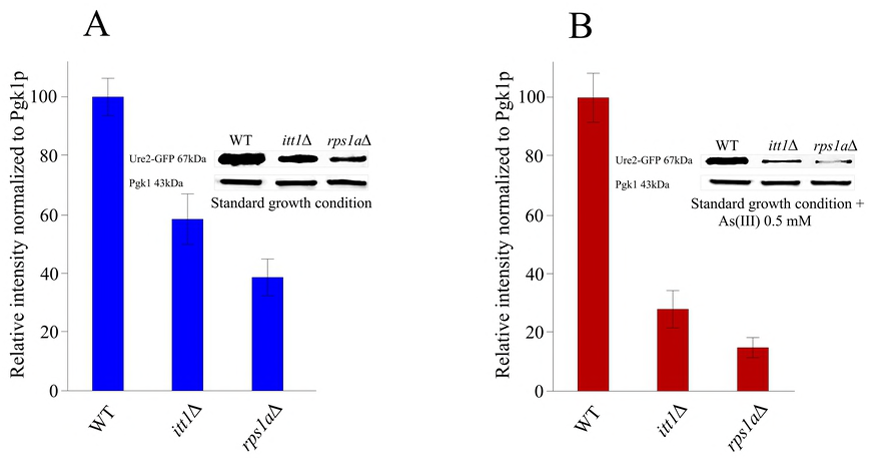
Genetic interaction (GI) analysis for *ITT1* and *RPS1A*. (A) Negative GIs for *ITT1* under standard growth conditions and the presence of As(III) or cycloheximide. In this case, deletion of a second gene along with *ITT1* forms an unexpected growth reduction. (B) Negative GIs for *RPS1A* under standard growth conditions and the presence of As(III) or cycloheximide as in (A). (C) Phenotypic suppression array (PSA) analysis using the over-expression of *ITT1*. In this way, overexpression for *ITT1* compensated for the sensitivity of gene deletions to As(III) (1.2 mM) or cyclohexamide (100 ng/ml). (D) PSA analysis using the overexpression of *RPS1A* as in (C). Each experiment was repeated three times and the interactions with 20% alteration or more in at least two screens were scored as positive. P-values were obtained from GeneMANIA [64].

Conditional GIs define an interesting type of GIs as they provide more compelling insight on the function of target genes under specific conditions [31], [58]. They represent the mosaic nature of gene function(s) which can change as a result of different external or internal factors. For example, although a number of genes are known to play a role in the DNA repair pathway, their expression is only regulated in response to the presence of DNA damage [29], [43]. We therefore investigated the negative GIs by assaying *ITT1* and *RPS1A* in the presence of a low concentration of As(III) (0.7 mM). Under this condition, we found both genes to form a new set of interactions with a series of genes that play a role in the regulation of translation (Figs 7a and b). This data suggests that in the presence of As(III), both genes appear to gain a new role in regulating the process of translation (p-value *ITT1* = 9.39e-5 and p-value *RPS1A* = 5.26e-10) (Figs 7a and b). Suppressor of ToM1 (*STM1*) is an example of a gene that formed new negative genetic interactions in the presence of As(III) with both *ITT1* and *RPS1A*. *STM1* codes for a protein that is required for optimal translation under nutrient stress [59], [60]. Enhancer of mRNA DeCapping *EDC1* and its paralog *EDC2* are other examples of the negative interactions gained by *RPS1A* under As(III) condition. EDC proteins directly bind to mRNA substrates and activate mRNA decapping. They also play a role in translation during stress conditions such as heat shock [61]. We also studied negative GIs in the presence of cycloheximide, which binds to the E-site of the 60S ribosomal subunit and interferes with deacetylated tRNA to inhibit general protein synthesis in the cell [62] (Figs 7a and b). Similar to our observations with As(III), in the presence of a mild concentration of cycloheximide (60 ng/ml), *ITT1* and *RPS1A* formed new negative GIs with a group of genes that are associated with the regulation of translation (p-value *ITT1* = 1.33e-6 and p-value *RPS1A*= 6.59e-9). Translation Initiation Factor 4A (*TIF2*) is an example of the gained interaction for both *ITT1* and *RPS1A* in response to cycloheximide. *TIF2* is a key player in translation initiation and holds a helicase activity [63]. Altogether, these observations are in agreement with the involvement of *ITT1* and *RPS1A* in regulating translation in response to stress.

To further study *ITT1* and *RPS1A*, we conducted phenotypic suppression array (PSA) analysis [31], [65] (Figs 7c and d) to assess the ability of overexpression of our target genes to reverse the defective phenotype (i.e. sensitivity) on a series of gene deletion strains under specific conditions. PSA analysis constitutes a more direct form of GI and can infer close functional relationships between interacting genes. In these cases, overexpression of one gene compensates the phenotypic adverse effect that is caused by the absence of another gene under certain conditions, such as stress caused by different chemicals [30], [31]. We observed that the overexpression of both *ITT1* and *RPS1A* reversed the sensitivity of a number of gene deletion strains to As(III) (1.2 mM) or cycloheximide (100 ng/ml) (Figs 7c and d). The majority of the newly identified gene interactors are involved in translation regulation further connecting the activity of *ITT1* and *RPS1A* to the regulation of translation. Interestingly, two of these genes, *GIS2* and *DOM34*, have reported to have IRES trans-acting factor (ITAF) activity [31], [66], [67]. The fact that the overexpression of *ITT1* and *RPS1A* can compensate for the absence of ITAFs, *GIS2* and *DOM34* provide further evidence connecting *ITT1* and *RPS1A* to IRES-mediated translation.

GIg Suppressor (*GIS2*) is a well studied translational activator for numerous IRES containing mRNAs [66], [67]. Duplication Of Multilocus region 34 (*DOM34*) is a protein that facilitates inactive ribosomal subunit dissociation to aid in translation restart [68], and is reported to play a role in IRES-mediated translation in yeast [31]. Our PSA analysis, not only supports a role in IRES-mediated translation pathway for both *ITT1* and *RPS1A*, but also suggests a possible systematic compensation between certain ITAFs which can be the subject of future studies. This also leads to the conclusion, that other interacting partners of *ITT1* and *RPS1A* in this experiment may play a role in IRES-mediated translation. Further studies are required to investigate these hypotheses. In agreement with our findings here, Sammons et al., [69] reported a physical interaction between Rpsa1p and Gis2p connecting the activity of these two proteins.

## References

1. Ferguson JE. The heavy elements: chemistry, environmental impact and health effects. 1990.

2. Tamás MJ, Martinoia E, editors. Molecular biology of metal homeostasis and detoxification. Berlin, Heidelberg, New York: Springer; 2006.

3. Goyer RA, Clarkson TW. Toxic effects of metals. Casarett & Doull’s Toxicology. The Basic Science of Poisons, Fifth Edition, Klaassen, CD [Ed]. McGraw-Hill Health Professions Division, ISBN. 1996;71054766.

4. Schwartz C, Gérard E, Perronnet K, Morel JL. Measurement of in situ phytoextraction of zinc by spontaneous metallophytes growing on a former smelter site. Science of the total environment. 2001 Nov 12;279(1-3):215–21.

5. Passariello B, Giuliano V, Quaresima S, Barbaro M, Caroli S, Forte G, Carelli G, Iavicoli Evaluation of the environmental contamination at an abandoned mining site. Microchemical Journal. 2002 Oct 1;73(1-2):245–50.

6. Chronopoulos J, Haidouti C, Chronopoulou-Sereli A, Massas I. Variations in plant and soil lead and cadmium content in urban parks in Athens, Greece. Science of the Total Environment. 1997 Mar 9;196(1):91–8.

7. Violante A, Cozzolino V, Perelomov L, Caporale AG, Pigna M. Mobility and bioavailability of heavy metals and metalloids in soil environments. Journal of soil science and plant nutrition. 2010 Jul;10(3):268–92.

8. Hernández RB, Nishita MI, Espósito BP, Scholz S, Michalke B. The role of chemical speciation, chemical fractionation and calcium disruption in manganese-induced developmental toxicity in zebrafish (Danio rerio) embryos. Journal of Trace Elements in Medicine and Biology. 2015 Oct 1;32:209–17.

9. Jaishankar M, Tseten T, Anbalagan N, Mathew BB, Beeregowda KN. Toxicity, mechanism and health effects of some heavy metals. Interdisciplinary toxicology. 2014 Jun 1;7(2):60–72.

10. Basu M, Bhattacharya S, Paul AK. Isolation and characterization of chromium-resistant bacteria from tannery effluents. Bulletin of environmental contamination and toxicology. 1997 Apr 1;58(4):535–42.

11. Choudhury P, Kumar R. Multidrug-and metal-resistant strains of Klebsiella pneumoniae isolated from Penaeus monodon of the coastal waters of deltaic Sundarban. Canadian journal of microbiology. 1998 Feb 1;44(2):186–9.

12. Castro-Silva MA, Lima AO, Gerchenski AV, Jaques DB, Rodrigues AL, Souza PL, Rörig LR. Heavy metal resistance of microorganisms isolated from coal mining environments of Santa Catarina. Brazilian Journal of Microbiology. 2003 Nov;34:45–7.

13. Otth L, Solís G, Wilson M, Fernández H. Susceptibility of Arcobacter butzleri to heavy metals. Brazilian journal of Microbiology. 2005 Sep;36(3):286–8.

14. Gadd GM. Transformation and mobilization of metals, metalloids, and radionuclides by microorganisms. Biophysico-chemical processes of heavy metals and metalloids in soil environments. Wiley, Hoboken. 2007 Nov 27:53–96.

15. Vadkertiová R, Sláviková E. Metal tolerance of yeasts isolated from water, soil and plant environments. Journal of basic microbiology. 2006 Apr 1;46(2):145–52.

16. Hosiner D, Gerber S, Lichtenberg-Frate H, Glaser W, Schüller C, Klipp E. Impact of acute metal stress in Saccharomyces cerevisiae. PLoS One. 2014 Jan 9;9(1):e83330.

17. Umland TC, Taylor KL, Rhee S, Wickner RB, Davies DR. The crystal structure of the nitrogen regulation fragment of the yeast prion protein Ure2p. Proceedings of the National Academy of Sciences. 2001 Feb 13;98(4):1459–64.

18. Bai M, Zhou JM, Perrett S. The yeast prion protein Ure2 shows glutathione peroxidase activity in both native and fibrillar forms. Journal of Biological Chemistry. 2004 Nov 26;279(48):50025–30.

19. Todorova TT, Kujumdzieva AV, Vuilleumier S. Non-enzymatic roles for the URE2 glutathione S-transferase in the response of Saccharomyces cerevisiae to arsenic. Archives of microbiology. 2010 Nov 1;192(11):909–18.

20. Rai R, Tate JJ, Cooper TG. Ure2, a prion precursor with homology to glutathione S-transferase, protects Saccharomyces cerevisiae cells from heavy metal ion and oxidant toxicity. Journal of Biological Chemistry. 2003 Apr 11;278(15):12826–33.

21. Urakov VN, Valouev IA, Lewitin EI, Paushkin SV, Kosorukov VS, Kushnirov VV, Smirnov VN, Ter-Avanesyan MD. Itt1p, a novel protein inhibiting translation termination in Saccharomyces cerevisiae. BMC molecular biology. 2001 Dec;2(1):9.

22. Swoboda RK, Broadbent ID, Bertram G, Budge S, Gooday GW, Gow NA, Brown AJ. Structure and regulation of a Candida albicans RP10 gene which encodes an immunogenic protein homologous to Saccharomyces cerevisiae ribosomal protein 10. Journal of bacteriology. 1995 Mar 1;177(5):1239–46.

23. Tong AH, Evangelista M, Parsons AB, Xu H, Bader GD, Pagé N, Robinson M, Raghibizadeh S, Hogue CW, Bussey H, Andrews B. Systematic genetic analysis with ordered arrays of yeast deletion mutants. Science. 2001 Dec 14;294(5550):2364–8.

24. Sopko R, Huang D, Preston N, Chua G, Papp B, Kafadar K, Snyder M, Oliver SG, Cyert M, Hughes TR, Boone C. Mapping pathways and phenotypes by systematic gene overexpression. Molecular cell. 2006 Feb 3;21(3):319–30.

25. Huh WK, Falvo JV, Gerke LC, Carroll AS, Howson RW, Weissman JS, O’shea EK. Global analysis of protein localization in budding yeast. Nature. 2003 Oct;425(6959):686.

26. Taylor RG, Walker DC, McInnes RR. E. coli host strains significantly affect the quality of small scale plasmid DNA preparations used for sequencing. Nucleic acids research. 1993 Apr 11;21(7):1677.

27. Komar AA, Lesnik T, Cullin C, Merrick WC, Trachsel H, Altmann M. Internal initiation drives the synthesis of Ure2 protein lacking the prion domain and affects [URE3] propagation in yeast cells. The EMBO journal. 2003 Mar 3;22(5):1199–209.

28. Winzeler EA, Shoemaker DD, Astromoff A, Liang H, Anderson K, Andre B, Bangham R, Benito R, Boeke JD, Bussey H, Chu AM. Functional characterization of the S. cerevisiae genome by gene deletion and parallel analysis. science. 1999 Aug 6;285(5429):901–6.

29. Omidi K, Hooshyar M, Jessulat M, Samanfar B, Sanders M, Burnside D, Pitre S, Schoenrock A, Xu J, Babu M, Golshani A. Phosphatase complex Pph3/Psy2 is involved in regulation of efficient non-homologous end-joining pathway in the yeast Saccharomyces cerevisiae. PloS one. 2014 Jan 31;9(1):e87248.

30. Alamgir M, Erukova V, Jessulat M, Azizi A, Golshani A. Chemical-genetic profile analysis of five inhibitory compounds in yeast. BMC chemical biology. 2010 Dec;10(1):6.

31. Samanfar B, Shostak K, Moteshareie H, Hajikarimlou M, Shaikho S, Omidi K, Hooshyar M, Burnside D, Márquez IG, Kazmirchuk T, Naing T. The sensitivity of the yeast, Saccharomyces cerevisiae, to acetic acid is influenced by DOM34 and RPL36A. PeerJ. 2017 Nov 14;5:e4037.

32. Samanfar B, Shostak K, Chalabian F, Wu Z, Alamgir M, Sunba N, Burnside D, Omidi K, Hooshyar M, Márquez IG, Jessulat M. A global investigation of gene deletion strains that affect premature stop codon bypass in yeast, Saccharomyces cerevisiae. Molecular Biosystems. 2014;10(4):916–24.

33. Szymanski EP, Kerscher O. Budding yeast protein extraction and purification for the study of function, interactions, and post-translational modifications. Journal of visualized experiments: JoVE. 2013(80).

34. Shaikho S, Dobson CC, Naing T, Samanfar B, Moteshareie H, Hajikarimloo M, Golshani A, Holcik M. Elevated levels of ribosomal proteins eL36 and eL42 control expression of Hsp90 in rhabdomyosarcoma. Translation. 2016 Jul 2;4(2):e1244395.

35. Esposito AM, Mateyak M, He D, Lewis M, Sasikumar AN, Hutton J, Copeland PR, Kinzy TG. Eukaryotic polyribosome profile analysis. Journal of visualized experiments: JoVE. 2010(40).

36. Faye MD, Graber TE, Holcik M. Assessment of selective mRNA translation in mammalian cells by polysome profiling. Journal of visualized experiments: JoVE. 2014(92).

37. Lou WP, Baser A, Klussmann S, Martin-Villalba A. In vivo interrogation of central nervous system translatome by polyribosome fractionation. Journal of visualized experiments: JoVE. 2014(86).

38. Sehgal A, Hughes BT, Espenshade PJ. Oxygen-dependent, alternative promoter controls translation of tco1+ in fission yeast. Nucleic acids research. 2008 Feb 14;36(6):2024–31.

39. Chassé H, Boulben S, Costache V, Cormier P, Morales J. Analysis of translation using polysome profiling. Nucleic acids research. 2016 Oct 7;45(3):e15-.

40. Tong AH, Lesage G, Bader GD, Ding H, Xu H, Xin X, Young J, Berriz GF, Brost RL, Chang M, Chen Y. Global mapping of the yeast genetic interaction network. science. 2004 Feb 6;303(5659):808–13.

41. Memarian N, Jessulat M, Alirezaie J, Mir-Rashed N, Xu J, Zareie M, Smith M, Golshani A. Colony size measurement of the yeast gene deletion strains for functional genomics. BMC bioinformatics. 2007 Dec;8(1):117.

42. Wagih O, Usaj M, Baryshnikova A, VanderSluis B, Kuzmin E, Costanzo M, Myers CL, Andrews BJ, Boone CM, Parts L. SGAtools: one-stop analysis and visualization of array-based genetic interaction screens. Nucleic acids research. 2013 May 15;41(W1):W591–6.

43. Tong AH, Boone C. 16 High-Throughput Strain Construction and Systematic Synthetic Lethal Screening in Saccharomycescerevisiae. Methods in Microbiology. 2007 Jan 1;36:369–707.

44. Omidi K, Jessulat M, Hooshyar M, Burnside D, Schoenrock A, Kazmirchuk T, Hajikarimlou M, Daniel M, Moteshareie H, Bhojoo U, Sanders M. Uncharacterized ORF HUR1 influences the efficiency of non-homologous end-joining repair in Saccharomyces cerevisiae. Gene. 2018 Jan 10;639:128–36.

45. Holcik M, Sonenberg N. Translational control in stress and apoptosis. Nature reviews Molecular cell biology. 2005 Apr;6(4):318.

46. Pelletier J, Sonenberg N. Internal initiation of translation of eukaryotic mRNA directed by a sequence derived from poliovirus RNA. Nature. 1988 Jul;334(6180):320.

47. Liwak U, Faye MD, Holcik M. Translation control in apoptosis. Exp Oncol. 2012 Sep;34(3):218–30.

48. Stoneley M, Willis AE. Cellular internal ribosome entry segments: structures, transacting factors and regulation of gene expression. Oncogene. 2004 Apr;23(18):3200.

49. Komar AA, Hatzoglou M. Cellular IRES-mediated translation: the war of ITAFs in pathophysiological states. Cell cycle. 2011 Jan 15;10(2):229–40.

50. Spriggs KA, Stoneley M, Bushell M, Willis AE. Re-programming of translation following cell stress allows IRES-mediated translation to predominate. Biology of the Cell. 2008 Jan 1;100(1):27–38.

51. Reineke LC, Cao Y, Baus D, Hossain NM, Merrick WC. Insights into the role of yeast eIF2A in IRES-mediated translation. PloS one. 2011 Sep 7;6(9):e24492.

52. Rieger R, Michaelis A, Green MM. Glossary of genetics and cytogenetics: classical and molecular. Springer Science & Business Media; 2012 Dec 6.

53. Szendro IG, Schenk MF, Franke J, Krug J, De Visser JA. Quantitative analyses of empirical fitness landscapes. Journal of Statistical Mechanics: Theory and Experiment. 2013 Jan 16;2013(01):P01005.

54. Kisly I, Gulay SP, Mäeorg U, Dinman JD, Remme J, Tamm T. The functional role of eL19 and eB12 intersubunit bridge in the eukaryotic ribosome. Journal of molecular biology. 2016 May 22;428(10):2203–16.

55. VanNice J, Gregory ST, Kamath D, O’Connor M. Alterations in ribosomal protein L19 that decrease the fidelity of translation. Biochimie. 2016 Sep 1;128:122–6.

56. Hannig EM, Hinnebusch AG. Molecular analysis of GCN3, a translational activator of GCN4: evidence for posttranslational control of GCN3 regulatory function. Molecular and cellular biology. 1988 Nov 1;8(11):4808–20.

57. Elsby R, Heiber JF, Reid P, Kimball SR, Pavitt GD, Barber GN. The alpha subunit of eukaryotic initiation factor 2B (eIF2B) is required for eIF2-mediated translational suppression of vesicular stomatitis virus. Journal of virology. 2011 Oct 1;85(19):9716–25.

58. Babu M, Díaz-Mejía JJ, Vlasblom J, Gagarinova A, Phanse S, Graham C, Yousif F, Ding H, Xiong X, Nazarians-Armavil A, Alamgir M. Genetic interaction maps in Escherichia coli reveal functional crosstalk among cell envelope biogenesis pathways. PLoS genetics. 2011 Nov 17;7(11):e1002377.

59. Utsugi T, Toh-e A, Kikuchi Y. A high dose of theSTM1 gene suppresses the temperature sensitivity of thetom1 andhtr1 mutants inSaccharomyces cerevisiae. Biochimica et Biophysica Acta (BBA)-Gene Structure and Expression. 1995 Sep 19;1263(3):285–8.

60. Nelson LD, Musso M, Van Dyke MW. The yeast STM1 gene encodes a purine motif triple helical DNA-binding protein. Journal of Biological Chemistry. 2000 Feb 25;275(8):5573–81.

61. Dunckley T, Tucker M, Parker R. Two related proteins, Edc1p and Edc2p, stimulate mRNA decapping in Saccharomyces cerevisiae. Genetics. 2001 Jan 1;157(1):27–37.

62. Schneider-Poetsch T, Ju J, Eyler DE, Dang Y, Bhat S, Merrick WC, Green R, Shen B, Liu JO. Inhibition of eukaryotic translation elongation by cycloheximide and lactimidomycin. Nature chemical biology. 2010 Mar;6(3):209.

63. Linder P, Slonimski PP. An essential yeast protein, encoded by duplicated genes TIF1 and TIF2 and homologous to the mammalian translation initiation factor eIF-4A, can suppress a mitochondrial missense mutation. Proceedings of the National Academy of Sciences. 1989 Apr 1;86(7):2286–90.

64. Warde-Farley D, Donaldson SL, Comes O, Zuberi K, Badrawi R, Chao P, Franz M, Grouios C, Kazi F, Lopes CT, Maitland A. The GeneMANIA prediction server: biological network integration for gene prioritization and predicting gene function. Nucleic acids research. 2010 Jun 21;38(suppl_2):W214–20.

65. Alamgir M, Eroukova V, Jessulat M, Xu J, Golshani A. Chemical-genetic profile analysis in yeast suggests that a previously uncharacterized open reading frame, YBR261C, affects protein synthesis. BMC genomics. 2008 Dec;9(1):583.

66. Scherrer T, Femmer C, Schiess R, Aebersold R, Gerber AP. Defining potentially conserved RNA regulons of homologous zinc-finger RNA-binding proteins. Genome biology. 2011 Jan;12(1):R3.

67. Sammons MA, Samir P, Link AJ. Saccharomyces cerevisiae Gis2 interacts with the translation machinery and is orthogonal to myotonic dystrophy type 2 protein ZNF9. Biochemical and biophysical research communications. 2011 Mar 4;406(1):13–9.

68. Davis L, Engebrecht J. Yeast dom34 mutants are defective in multiple developmental pathways and exhibit decreased levels of polyribosomes. Genetics. 1998 May 1;149(1):45–56.

69. Sammons MA, Samir P, Link AJ. Saccharomyces cerevisiae Gis2 interacts with the translation machinery and is orthogonal to myotonic dystrophy type 2 protein ZNF9. Biochemical and biophysical research communications. 2011 Mar 4;406(1):13–9.

